# Impact of Model Order Choice on the Results of Parallel Independent Component Analysis

**DOI:** 10.1101/2020.08.31.276196

**Authors:** D. M. Jensen, E. Zendehrouh, J. Liu, V. D. Calhoun, J. A. Turner

## Abstract

Parallel independent component analysis (pICA) is a data-driven method that identifies the maximally independent components of multiple imaging modalities while simultaneously investigating the strength of their correlations. Researchers using pICA are given the option to use the suggested model order calculated by the minimum descriptive length (MDL) algorithm, or they can choose their own model order. To date, there are no suggested guidelines for this choice. To test the sensitivity of pICA to the selection of model order, we applied it to a well-researched brain disorder, schizophrenia, looking at the correlations between patterns of grey matter volume (GM) volume and white matter integrity, measured using fractional anisotropy (FA). We varied model orders from low to high, and tested the sensitivity to disorder effects (cases vs controls), similarity of spatial maps identified across model orders, consolidation or distribution effects related to model order selection, and the performance of the minimum descriptive length (MDL) algorithm. The pICA results (multimodal analysis) were also compared to the ICA (unimodal analysis) for each imaging modality. Across model orders, there was consistent sensitivity to disorder effects, and clustered patterns of spatial maps for both the GM and FA reflecting those differences. The MDL-estimated model order captured the majority, but not all, of the spatial patterns present in the GM and FA. There was not the expected consolidation of spatial maps at lower model orders, nor the distribution of spatial maps at higher model orders. The spatial patterns identified in the ICA closely resemble those found in the pICA, although lacking the benefit of the optimization algorithm, were not as highly correlated. This offers some insight and guidance for researchers interested in using pICA with regard to selecting model order for their particular analysis of multiple imaging modalities.

## I. Introduction

Multimodal analysis, when applied to magnetic resonance imaging (MRI), offers an opportunity to take the understanding of brain function and dysfunction a step further by fusing different modalities of imaging data, allowing deeper insights into the relationships between already well-developed bodies of knowledge [1]. As yet, these types of analysis are underutilized and deserve more exploration. A multimodal analysis of two different image modalities, for example, will search for patterns of interactions, revealing relationships that a single modality cannot bring to light.

There are two main types of multimodal analysis: model-driven and data-driven. Model-driven multimodal analysis uses an a priori hypothesis, requiring a specific question based on previous knowledge of the problem, such as a general linear model or dynamic causal modeling. In contrast, data-driven multimodal analysis is a relatively hypothesis-free method of investigating neuroimaging that allows exploration of the data without needing a hypothesis [2]. Parallel independent component analysis (pICA) can be a hybrid of the two, a semi-blind method that facilitates a data-driven whole brain exploration of the relationships between the two imaging modalities, while allowing some input from the researchers into the algorithm, including selection of the model order used. When the MDL is used to estimate the number of components from the data, pICA is then a data-driven analysis. [3].

pICA takes an already powerful tool, independent component analysis (ICA), and uses it to identify the maximally independent components of multiple modalities while simultaneously investigating their correlations [4]. Used in conjunction with ICASSO [32], an algorithm designed specifically to improve the stability of estimated components for neuroimaging data, pICA can be used to both identify and quantify the relationships between the features, or spatial patterns in the brain images. This emphasis on the pattern of interactions also makes it more robust to noise, a non-trivial consideration when dealing with neuroimaging [2]. Individual variance between subjects is reflected in the loading coefficients of the features.

Figure 1 outlines the pICA process. Once the components for each modality have been identified, the bridge between the two data sets is the constraint, which determines the optimal interconnection between them. Two safeguards are used to prevent over/underestimating the existing interrelationships: the dynamic updating of the constrained interconnections, allowing them to vary between iterations, and the adaptive learning rates of the terms within the cost function, which are updated in parallel. For a more complete description of the algorithm, see [24].

**Figure 1:**
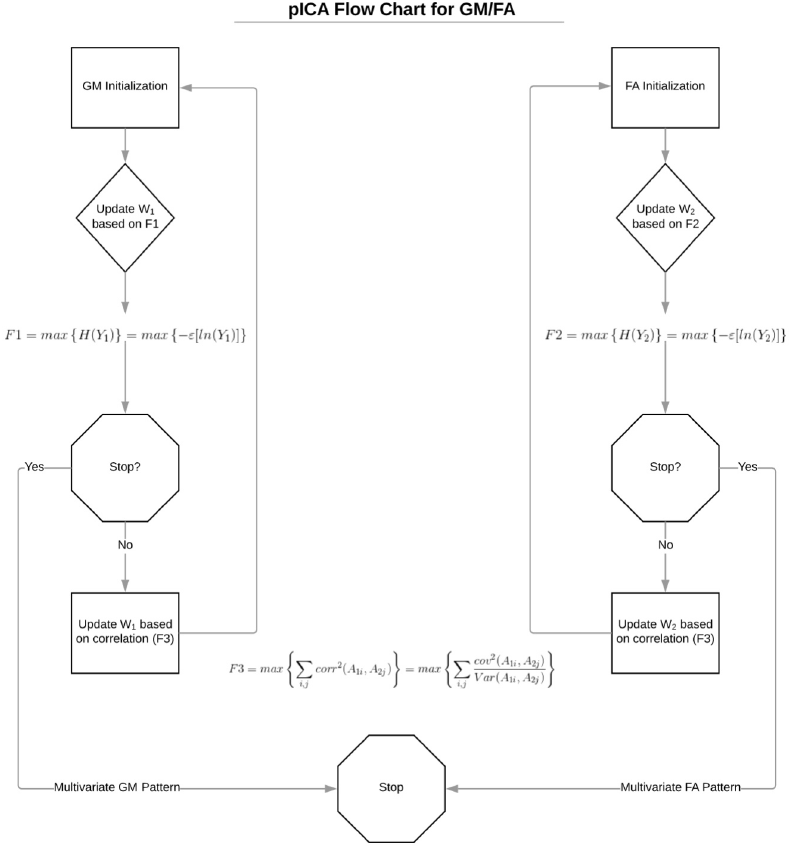
The parallel independent component analysis iteration optimization procedure, when complete, finds the link between two multivariate patterns and calculates their correlation, creating the correlated pairs of FA/GM components [3].

When using ICA, a key issue for researchers is determining the amount of data-reduction that should be applied to the images before the analysis, reflected in the choice of component numbers. A default method, the minimum description length (MDL) algorithm, should suggest an optimal number of components and is provided within the software package. This algorithm has proven useful in decomposing fMRI data (ICA), but to date has not been explicitly tested on sMRI or DTI, nor has its performance when used with fusion analysis been examined. [19] We undertook to test the effect of model order, or number of components, on the sensitivity to case/control differences and the identified spatial patterns. We wanted to see the impact, if any, model order choice had on the consolidation and/or distribution of the networks with significant group differences. We also wanted to compare the performance of the MDL to other model orders with regard to its ability to capture the spatial maps identified by the pICA. We ran successive iterations of the pICA using varied numbers of components and constraints to discover how many pairs would be significantly correlated, how many would show significant group differences, and how similar the spatial maps identified were across differing model orders. We also ran individual ICAs on both the image modalities using source-based morphometry (SBM) at varied model orders to test the similarity of those spatial maps that had significant group differences and to compare those to the ones found in the pICA at the MDL suggested model order.

To do this effectively, we chose a well-studied disorder, schizophrenia, and used a well-researched data set (COBRE) [25]. While few have explored the relationship between the grey and white matter changes that characterize this disorder, the diffuse, diverse, and wide-spread changes in both neural types are well-documented, and previous unimodal research establishes a global association between the reduction in grey matter and white matter integrity loss [1,5,6,7]. There have been numerous multimodal analyses investigating various aspects of schizophrenia, including a study using joint ICA (jICA) that identified group differences within joint sources of grey and white matter volume between healthy controls (HC) and patients with schizophrenia (SZ) [8]. Few have applied pICA to the grey matter volume estimates (GM) from the sMRI images and the fractional anisotropy (FA) measurements from the DTI images to probe the relationship between the volume of the grey matter and the integrity of the white matter in HC compared to SZ.

## II. Methods

### Subjects

The images were collected from 157 subjects, 82 healthy controls (HC) and 75 subjects with schizophrenia (SZ) as part of the COBRE dataset [25]. Two subjects with schizophrenia were removed for poor imaging data. The final total was 155, with the HC group having 62 males and 20 females, and the SZ group having 59 males and 15 females. Both groups ranged from 18-65 years in age. The Structured Clinical Interview for DSM Disorders (SCID) was used to gather diagnostic information and subjects were excluded if they had history of substance abuse or dependence within the last 12 months, severe head trauma with more than 5 minutes loss of consciousness, neurological disorders, or severe cognitive impairment. All subjects provided informed consent prior to the study. A Welch’s two sample t-test and a Pearson’s chi-squared test were used to test for group differences in age and gender respectively using R version 3.5.0.

### Image Collection

The data were collected on a Siemens 3T Trio TIM scanner at the Mind Research Network, Albuquerque, NM.

The T1-weighted images for GM were collected in the sagittal plane, interleaved, multi-slice mode in a single shot with these parameters: TR/TE/TI = 2530/[1.64, 3.5, 5.36, 7.22, 9.08]/900 ms, flip angle = 7 degrees, FOV = 256×256 mm, matrix 256×256×176, voxel size =1×1×1 mm, number of echos = 5, pixel bandwidth = 650 Hz, total scan time = 6 min.

The dMRI DTI images for FA were collected using 30 gradient directions and 5 b=0, for a total of 72 slices with a slice thickness of 2mm (isotroptic resolution of 2×2×2 mm). FOV=256×256 mm, TR/TE=9000 ms/84 ms, encoded A-P. Sequence bandwidth was 1562 Hz/Px and echo spacing was 0.72 ms with an EPI factor of 128. For more information, see Aine et al [30]

### Image Processing

#### dMRI to FA

An FSL v5.0.10 pipeline was used to preprocess the DTI data [9]. A quality control of the DTI images was done using DTIPrep to ensure that a minimum of 25 gradient directions for each subject were free of artifacts [31]. Eddy current correction for gradient distortions and head motion were applied to the diffusion-weighted images [10], after which a brain extraction tool (BET) was used to remove non-brain tissue from the image [11]. A diffusion tensor model was fitted to each voxel with DTIFIT [12], creating the fractional anisotropy images. All subjects’ FA data were then aligned into a common space using the nonlinear registration tool FNIRT [27,28], which uses a b-spline representation of the registration warp field [29]. Leaving the FA unsmoothed and in 1×1×1 MNI152 resolution eliminated spurious results due to partial voluming.

#### sMRI to GM

The T1-weighted sMRI images were reoriented and registered to the MNI152 template and resampled to 2mm x 2mm x 2mm. Using DARTEL in SPM12, a high-dimensional normalization pipeline [16], the non-brain tissues were stripped and the grey matter, white matter, and cerebral spinal fluid were segmented, leaving normalized, modulated, Jacobian-scaled grey matter images that were then smoothed by an 8mm x 8mm x 8mm Gaussian kernel.

#### pICA

Parallel ICA was performed using the Fusion ICA Toolbox (FITv2.0a) run in Matlab R2017b. See Figure 1 for algorithm details. For the initial iteration, principle components for each modality were estimated using a minimum description length in the FIT software (4 FA components and 52 GM components when estimated separately, 12 when combined) [17]. The descending trend of entropy was allowed to be −0.001 maximally. ICASSO software was used to ensure cluster stability by retesting each FastICA 10 times.

Ten different pICA were run with different model orders, i.e. with the number of components ranging from 11 to 39 for both features. Each model order was constrained in the number of correlations to roughly half the component number. For each pICA, each subject’s data are broken down into common GM and FA spatial patterns (components), and the loading coefficients for these components.

We identified the case/control differences in the loading coefficients for the correlated pairs of GM/FA components in each model order using a two-sample t-test. Z-scores of the spatial patterns were thresholded at |z| > 3 to identify component clusters.

### Comparisons Across Model Order

Comparisons across model order were made using FSL to examine the consistency of the FA and GM components identified with significant group differences. A correlation analysis was done first, using the absolute values of the FA and GM component images and then correlated across model numbers using *fslcc*. The results were compiled in a heatmap using R version 3.5.0 and clustered to reveal repeating patterns of components across different model orders. A Jaccard Index of similarity was calculated next for each modality to explore the possibility of subsetting (consolidation/distribution) of spatial patterns across the model orders. *Fslmaths* was used to create images that calculated the ratio between the intersection of two components and their union. A heatmap of the similarity index for each modality was created in R.

### Correlation analysis across model orders using the source-based morphology (SBM)

SBM is the ICA of each modality in isolation using the gift toolbox (SBM v1.0b). For both FA and GM, 10 different model orders were tested using parameters identical to the pICA process. A correlation matrix and resulting heatmap for each modality was calculated using R, focusing on the spatial maps with significant group differences as identified using the gift ANOVA toolbox. A correlation between the FA and GM spatial maps with significant group differences from the MDL recommended number of components for the pICA (12 components) was done in R to identify relationships between the two modalities to be compared to the pICA correlated spatial maps.

## III. Results

### pICA Results

All 10 pICA models were successful at finding significantly correlated pairs of components (multiple comparison correction 0.05/number of components per model order, see Supplementary Table 1), as well as component pairs with significant group differences. See Table 1.

**Table 1:** Summary of the pICA iterations: Listed here are the model orders investigated, how many correlation constraints per model order, the number of significantly correlated component pairs found (p corrected for multiple comparisons for each model order investigated), how many significantly correlated component pairs had significant group differences (two-sample t-test), and the highest correlation value for the significant component pairs with significant group differences found by the pICA.

| Model Order | Constraints | Significantly Correlated Pairs | Pairs with Significant Case vs Control Difference | Highest r-value of Significantly Correlated Pairs |
| --- | --- | --- | --- | --- |
| 11x11 | 5 | 5 | 2 | 0.73 |
| 12x12 | 6 | 6 | 3 | 0.61 |
| 15x15 | 8 | 8 | 4 | 0.81 |
| 17x17 | 9 | 9 | 3 | 0.86 |
| 20x20 | 10 | 10 | 1 | 0.72 |
| 23x23 | 12 | 12 | 5 | 0.78 |
| 28x28 | 14 | 14 | 3 | 0.75 |
| 33x33 | 17 | 17 | 2 | 0.54 |
| 36x36 | 18 | 18 | 5 | 0.82 |
| 39x39 | 19 | 17 | 1 | 0.42 |

### Comparisons Across pICA Model Order Results

In figures 2a and 2b, we see the frequency of correlation coefficients by model order between the spatial maps of the FA and GM respectively. The majority of the model orders for FA had a mean of correlation coefficients around 0.60, showing a consistent relationship between the spatial maps across model orders. The frequency distribution for the correlation of GM spatial maps showed some inconsistency in their relationships across model orders. The lower model orders (11×11, 12×12, and 15×15) had a mean of correlation coefficients of 0.40, but in the higher models orders (17×17 - 39×39), the mean of the correlation coefficients steadily drops from 0.40 to 0.35, showing a drop in consistency in the relationship between the spatial maps as the model order increases, as well as a lower consistency in the GM spatial maps across model orders compared to the FA spatial maps.

**Figure 2a and b:**
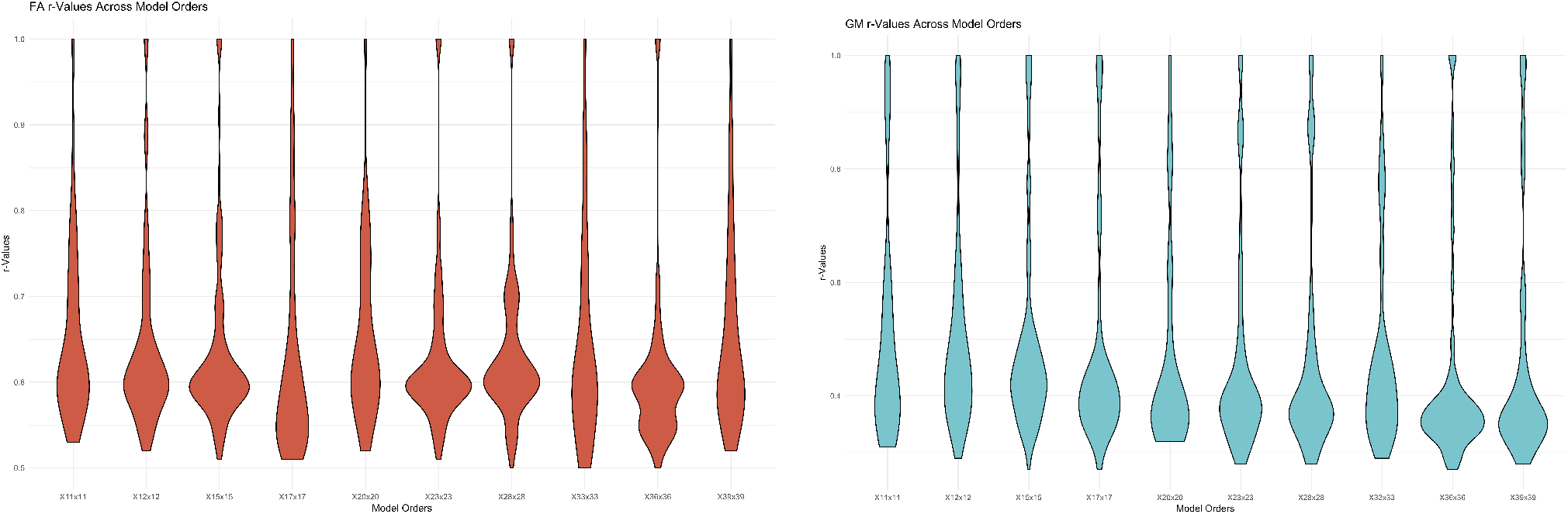
FA and GM Violin plots respectively, showing the frequency of the correlation indexes per model order.

Figures 3 and 4 are heat maps showing the results of the correlation analysis between the spatial maps with group differences of the pICA components for FA and GM across the 10 different model orders. The GM results showed more distinct clusters of related spatial maps than the FA. The 3 spatial maps of the mdl recommended model order represented the main clusters of spatial maps found across model orders in the FA results, but did not represent all the spatial map clusterings from the GM.

**Figure 3:**
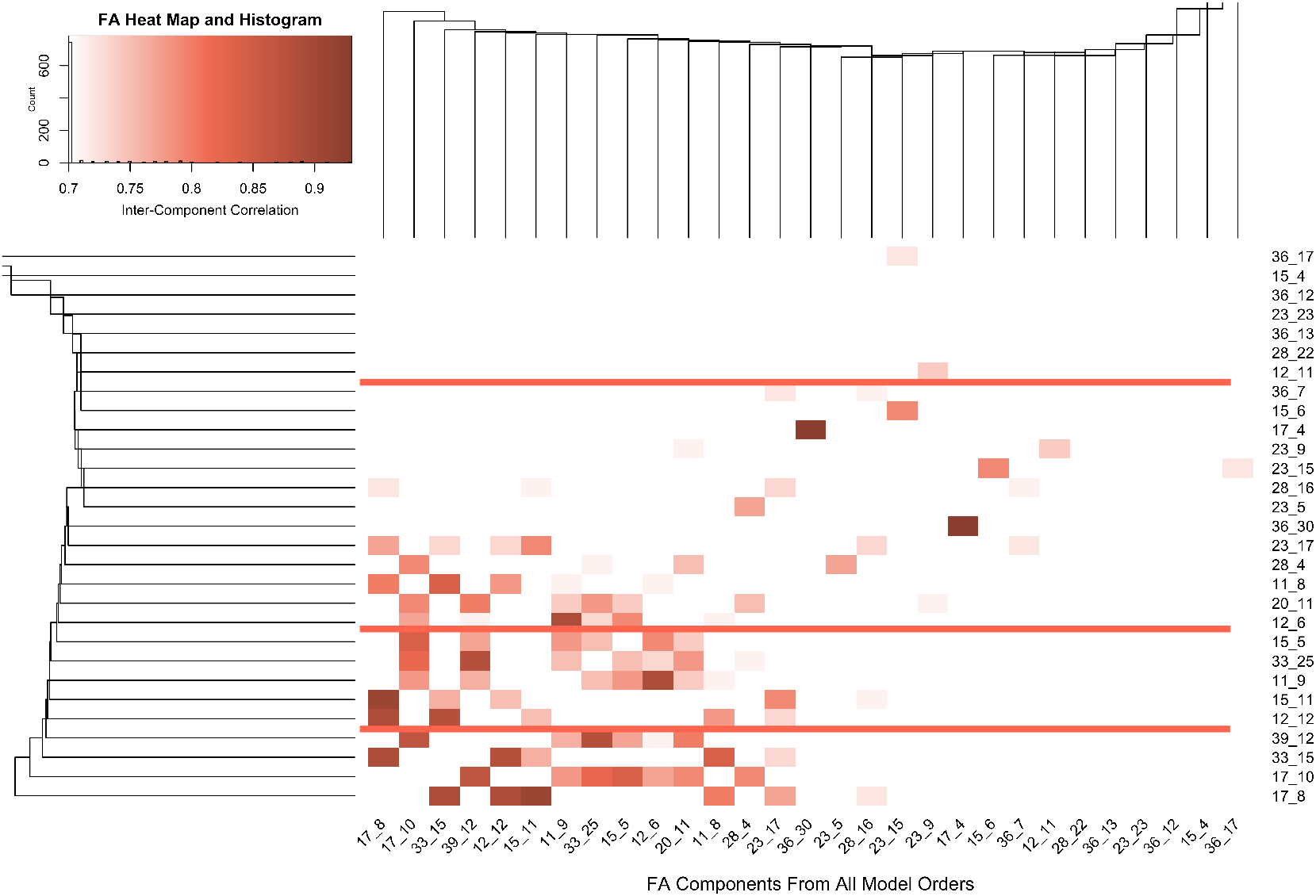
Heat map and hierarchical cluster analysis of FA spatial maps with group differences across model orders, thresholded at 0.7. Each axis lists the pICA model order number, and the component number. (i.e. 12_6 is from model order 12×12 and it’s the 6^th^ FA component identified in the 12×12 model order). The branches of the dendrogram represent the dissimilarities between the clusters using their squared Euclidian distances. The majority of the highest correlated pairs (r > 0.90) are in the bottom left corner of the largest cluster. The red underscore highlights the relationship of the MDL recommended 12×12 model order.

**Figure 4:**
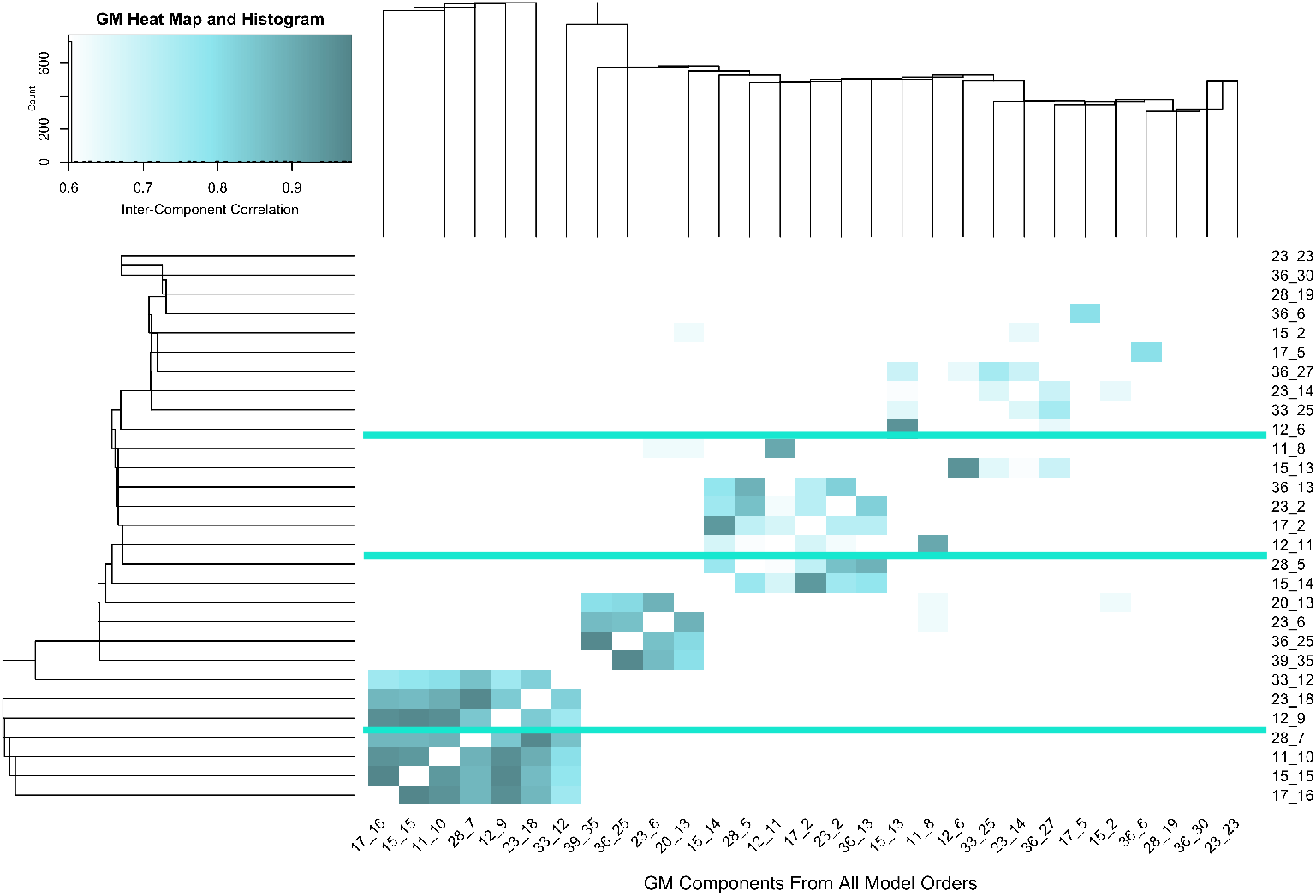
Heat map and hierarchical cluster analysis of GM spatial maps with group differences across model orders, thresholded at 0.6. Each axis lists the pICA model order number and the component number. (i.e. 36_25 is from model order 36×36 and it’s the 25^th^ GM component from the 36×36 model order). The branches of the dendrogram represent the dissimilarities between the clusters using their squared Euclidian distances. The majority of the highest correlated pairs (r > 0.90) are in the bottom left corner of the largest cluster. The green underscore highlights the relationship of the MDL recommended 12×12 model order.

Figures 5 and 6 show a heat map of the results of the Jaccard index for the FA and GM respectively, thresholded at 0.0 and higher, highlighting the effect of model order on the amount of subsetting between the spatial maps. Overall, the Jaccard index for FA was lower than GM, showing a lesser degree consolidation (overlap) of the spatial maps at the lower model orders, and a high degree of distribution (very little overlap) of the spatial maps at the higher model orders. The GM results show a much greater degree of consolidation of the spatial maps overall, with more distribution of the spatial maps at the higher model orders.

**Figure 5:**
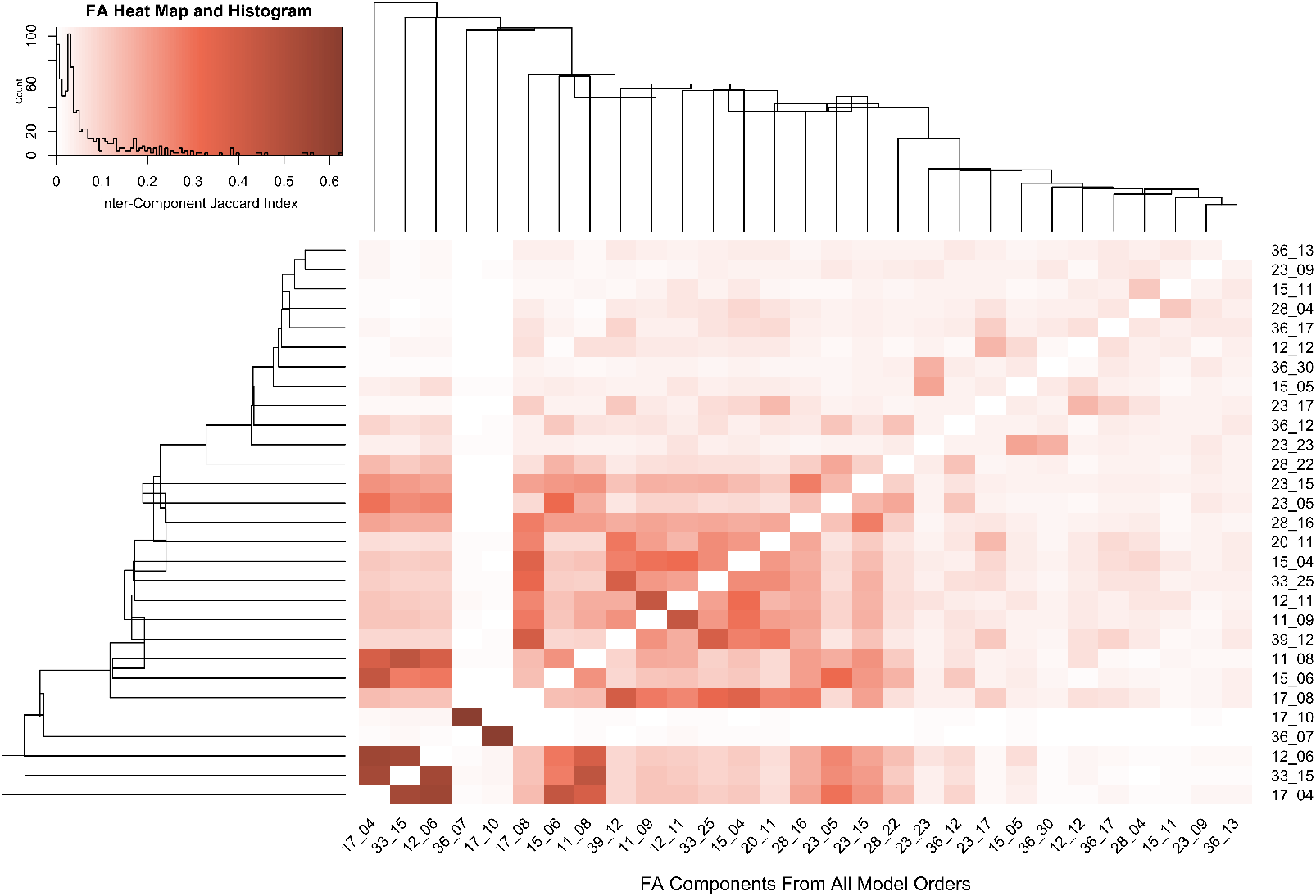
Heatmap of the FA Jaccard Index of Similarity, looking at subsetting between the spatial maps with group differences across model orders. Each axis lists the pICA model order number, and the component number. (i.e. 12_6 is from model order 12×12 and it’s the 6^th^ FA component identified in the 12×12 model order). The branches of the dendrogram represent the dissimilarities between the clusters using their squared Euclidian distances. The higher the Jaccard Index, the more shared information between the spatial maps (overlap).

**Figure 6:**
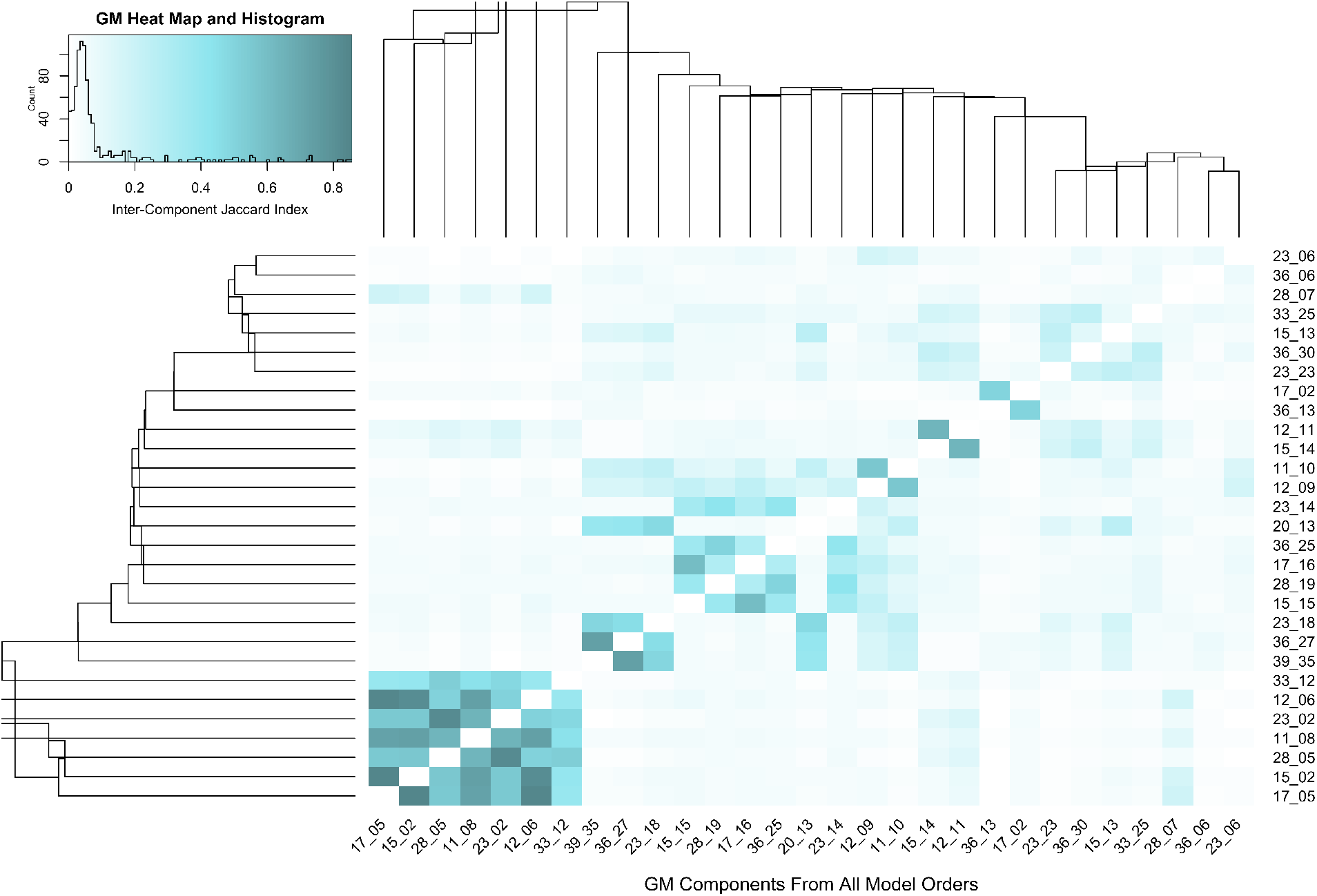
Heatmap of the GM Jaccard Index of Similarity, looking at subsetting between the spatial maps with group differences across model orders. Each axis lists the pICA model order number, and the component number. (i.e. 36_25 is from model order 36×36 and it’s the 25^th^ GM component from the 36×36 model order). The branches of the dendrogram represent the dissimilarities between the clusters using their squared Euclidian distances. The higher the Jaccard Index, the more shared information between the spatial maps (overlap).

### Comparisons Across SBM Model Order Results

Figures 7 and 8 show the a heat map of the results of the correlation analysis of the SBM per modality. The GM showed more distinct clustering of spatial maps compared to the FA results. Using the pICA MDL recommendation of 12 components, the simplistic correlation analysis of the FA loading coefficients and the GM loading coefficients from the SBM returned 3 pairs of correlated FA and GM spatial maps with significant group differences.

**Figure 7:**
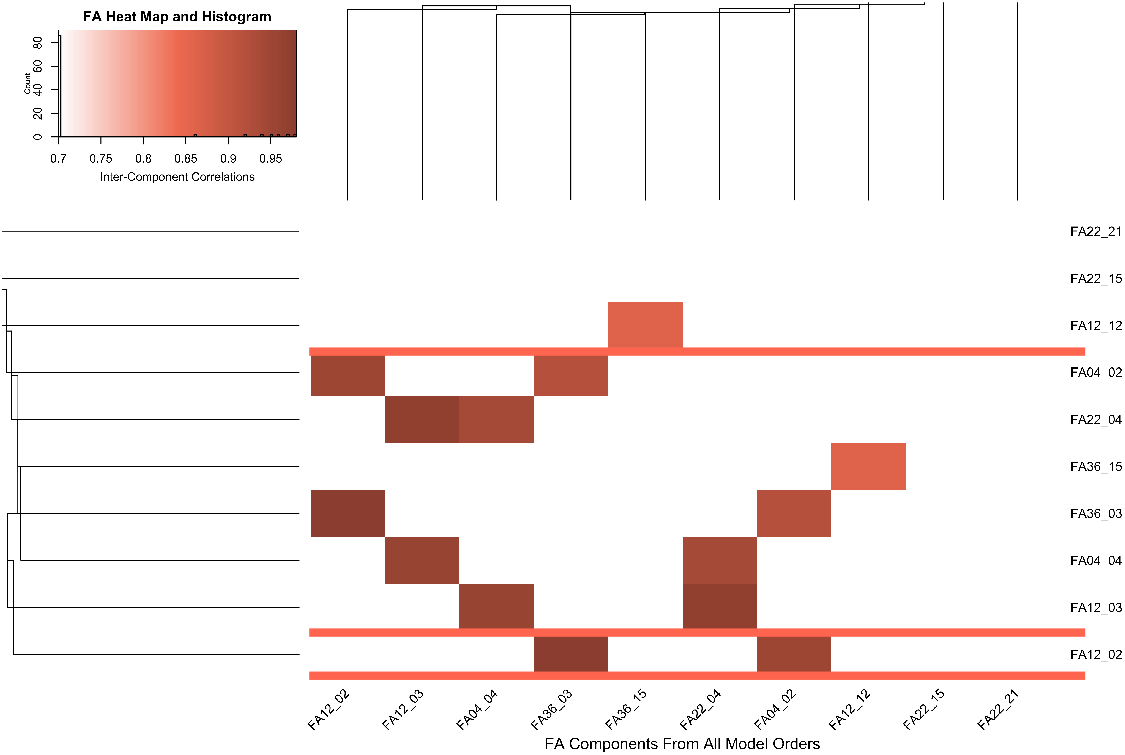
Heat map and hierarchical cluster analysis of the SBM (unimodal) FA spatial maps with group differences across model orders, thresholded at 0.7. Each axis lists the SBM model order number, and the component number. (i.e. FA12_02 is from model order 12×12 and it’s the 2nd FA component identified in the 12×12 model order). The branches of the dendrogram represent the dissimilarities between the clusters using their squared Euclidian distances. The majority of the highest correlated pairs (r > 0.90) are in the bottom left corner of the largest cluster. The red underscore highlights the relationship of the MDL recommended 12×12 model order.

**Figure 8:**
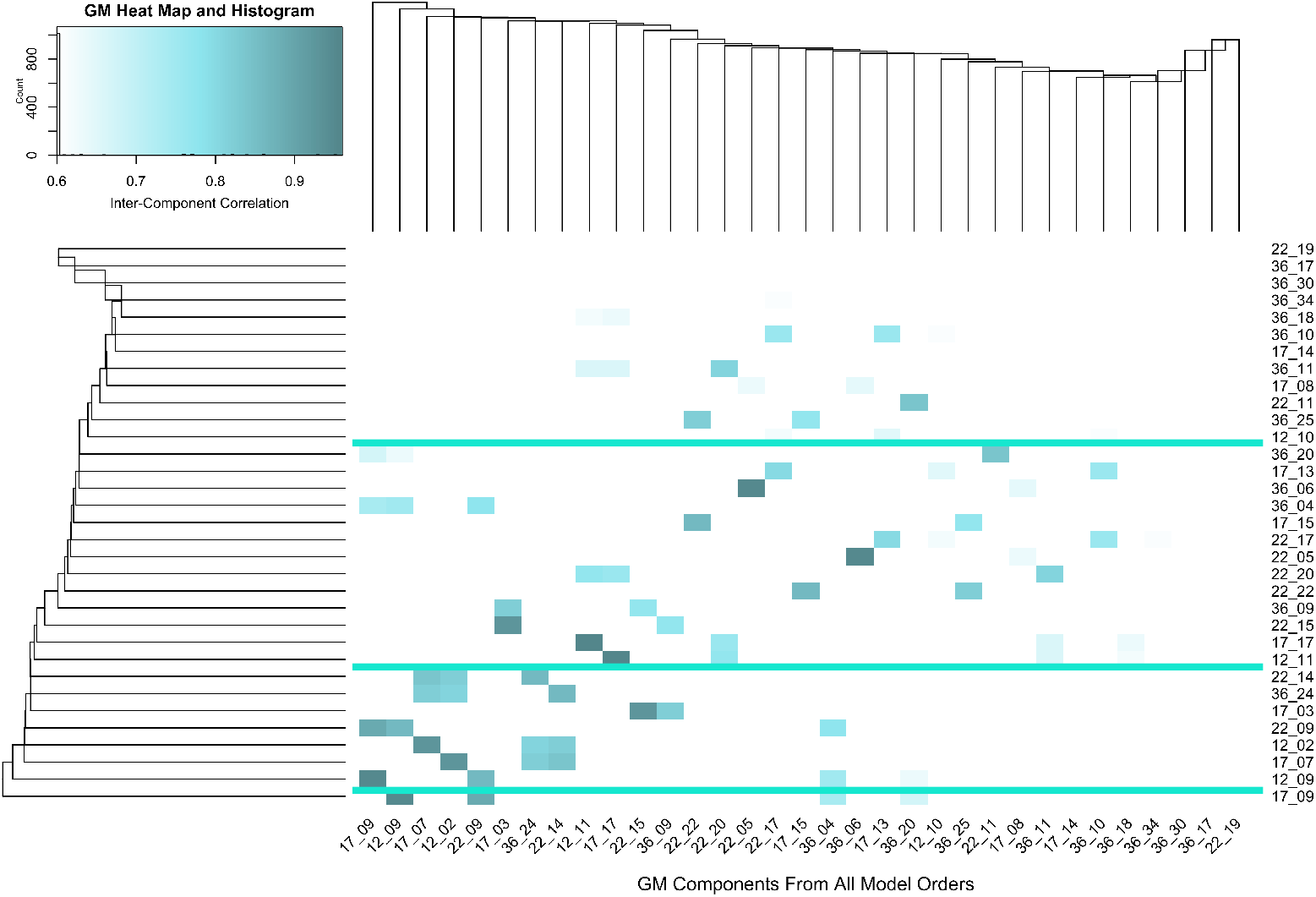
Heat map and hierarchical cluster analysis of the SBM (unimodal) GM spatial maps with group differences across model orders, thresholded at 0.6. Each axis lists the pICA model order number and the component number. (i.e. 36_06 is from model order 36×36 and it’s the 6t^h^ GM component from the 36×36 model order). The branches of the dendrogram represent the dissimilarities between the clusters using their squared Euclidian distances. The majority of the highest correlated pairs (r > 0.90) are in the bottom left corner of the largest cluster. The green underscore highlights the relationship of the MDL recommended 12×12 model order.

Figures 9 compares the spatial maps returned by the SBM and the pICA analysis. There is a high degree of similarity between the spatial maps from both analyses, although the pICA results had higher correlation coefficients than those returned by the simple SBM correlation analysis.

**Figure 9:**
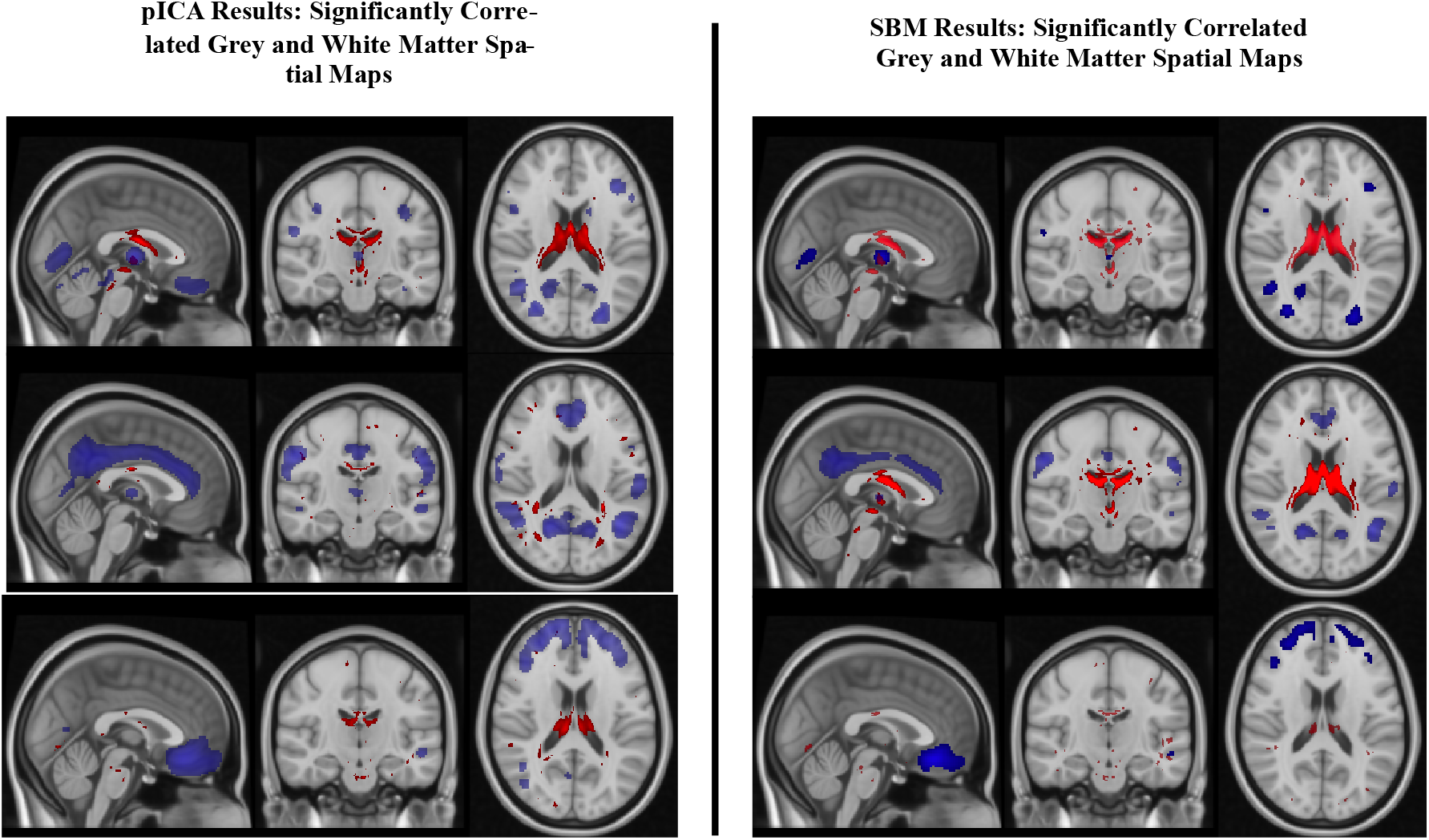
Comparisons of the 12×12 model order results, pICA vs SBM: Side-by-side comparison of the significantly correlated GM and WM spatial maps with significant group different found using the pICA and the simple correlation run in R using the SBM results. Both identified very similar brain regions, although the pICA optimization returned higher correlation coefficients.

## IV. Discussion

A fundamental assumption of ICA is the independent and identically distributed nature of the data. MRI images violate that assumption due to the inherent dependence of one voxel to another as well as the further reduction of independence once Gaussian smoothing is applied to the images. [17] This can be seen in the difference in the degree of relationship between the spatial maps across model orders for the FA and GM in both the pICA and the SBM results. The FA was less independent (had a higher degree of relatedness) across the model orders, while the GM displayed distinct clusters of independent spatial maps across different model orders. The MDL, designed to correct for the misestimation of model order caused by sample dependence, did capture most of the spatial patterns returned by the pICA and SBM in the GM, it did not represent them all. The FA spatial maps were more accurately represented in the MDL recommended 12×12 model order. Further experimentation with different Gaussian kernels to smooth both the GM and FA could explain the varied performance of the MDL algorithm and improve its accuracy.

Both the FA and the GM showed the expected consolidation of spatial maps at the lower model orders (higher degree of similarity/overlap) and the distribution of the spatial maps at higher model orders (lower degree of similarity/overlap), although the FA had a lower degree of similarity overall compared to the GM. Again, differences in smoothing could account for this and should be explored further.

The unimodal ICAs of FA and GM identified spatial maps whose loading coefficients showed significant group differences that identified similar brain regions as those found in the pICA. The correlation results returned from the analysis run in R were lower, highlighting the benefit of the optimization algorithm within the pICA. While it is fairly simple to run a simplified correlation analysis between two modalities, it would be impossible to do this with three or more modalities. This is one of the main benefits of the pICA algorithm, that it not only optimizes the correlation between modalities, but that it allows researchers to compare three or more modalities. [24]

The pICA was consistently sensitive to case vs controls differences across model orders. There was no indication that lower and higher model orders were more or less correlated with one another.

## V. Conclusion

Researchers considering pICA as an analysis tool combining grey matter volume and fractional anisotropy images should use the least amount of smoothing necessary for their particular image modality to maximize the likelihood that the MDL suggested model order is as reflective of all the spatial maps within their data. The pICA was otherwise robust and consistently identified correlated group differences between SZ and HC.

